# Ecological memory mitigates negative impacts of disturbance on biomass production in benthic diatom metacommunities

**DOI:** 10.1101/2020.07.23.217448

**Authors:** Friederike G. Engel, Birte Matthiessen, Rosyta Andriana, Britas Klemens Eriksson

**Author notes:** Corresponding author: Friederike G. Engel, Department of Biology, University of Florida, 876 Newell Dr, Gainesville, FL 32611, USA.

## Abstract

Disturbance events to coastal habitats such as extreme heat events, storms, or floods have increased in magnitude and frequency in recent years due to anthropogenic climate change and the destruction of habitats. These events constitute a major threat to many ecological communities and global biodiversity. Disturbance history influences ecosystem response to novel disturbances such that communities that have previously been exposed to disturbances should be more resilient to new disturbances compared to previously sheltered communities. This principle is defined as ecological memory. Resilience should also increase with access to a larger species pool, because a larger species pool increases species and response diversity of a community. One possibility of increasing the local species pool is connectivity via adequate dispersal between habitat patches with different species compositions in metacommunities. In a laboratory experiment, we exposed benthic diatom communities of different origin to a mechanical disturbance, simulated dispersal in half of the communities, and measured their chlorophyll *a* concentration over time. The local diatom communities originated from different locations on an intertidal flat that varied in hydrodynamic exposure history. Hydrodynamic exposure disturbs the sediment, and thereby determines sediment properties and the composition of intertidal diatom communities. In the experiment, disturbance negatively affected chlorophyll *a* concentration across all treatments. However, the response to disturbance depended on the ecological memory of the communities; the more exposed areas the communities originated from, the less negative was the effect of the mechanical disturbance. Interestingly, dispersal did not mitigate the negative impacts of disturbance in any of the communities. Our results highlight the importance of ecological memory for ecosystem functioning and demonstrate the limitations of patch connectivity to alleviate the impacts of disturbance events in metacommunities.

## Introduction

Global climate change and habitat destruction have altered many ecosystems which poses an urgent threat to many ecological communities and thus to global biodiversity (IPCC 2014). In addition to increased average global temperatures, the severity and frequency of extreme weather events such as storms and floods are expected to increase in the future (Harley et al. 2006, IPCC 2014). These extreme events will severely affect coastal areas, including the North Sea coast (Beniston et al. 2007), where they disturb and redistribute surface sediments on intertidal flats (Bartholomä et al. 2009). Increased sediment dynamics caused by storms and floods will most likely affect intertidal production negatively, because sediment erosion is the main abiotic constraint for autotrophic organisms living in and on surface sediments (de Jonge and van Beusekom 1995, Donadi et al. 2013a). Resilience in the face of disturbances is crucial for the survival of ecological communities and the maintenance of ecological functions in our fast-changing world (Oliver et al. 2015, König et al. 2019).

Species diversity affects ecosystem functioning (Hooper et al. 2005, Cardinale et al. 2012, Gonzalez et al. 2020) and influences ecosystem responses to disturbances by determining the system’s response diversity and resilience (Elmqvist et al. 2003, Mori et al. 2013). Higher response diversity increases the resilience of communities and thus assures the maintenance of ecosystem functioning during and after disturbance events (Elmqvist et al. 2003, Mori et al. 2013, Carrara et al. 2015, Oliver et al. 2015). Species and response diversity are increased in communities with access to a larger, regional species pool, compared to local isolated communities, if the different local communities have varying species compositions (Altermatt et al. 2011, Cosentino et al. 2011). Individual local communities that are connected via dispersal (i.e. the passive or active movement of individuals from one local patch to another) form metacommunities (Gilpin and Hanski 1991, Wilson 1992, Leibold et al. 2004, Holyoak et al. 2005). Increased species and response diversity in metacommunities are mainly caused by the sampling effect (Loreau and Hector 2001) and the spatial insurance effect (Loreau et al. 2003, Leibold and Chase 2018). Both these principles are based in the theory that access to a regional species pool with diverse traits increases the probability of local patch colonization by superior species that can maximize ecosystem functioning. In addition, mass effects (Mouquet and Loreau 2003) that lead to the constant replenishment of biomass from the regional species pool, and thus supply regional dominant species to local patches, can aid in the resilience of communities.

Previous states and experiences can affect future responses of communities, a process coined as “ecological memory” (Padisak 1992, Ogle et al. 2015). Ecological memory can manifest in different ways including the retention of certain physiological, behavioral, morphological, molecular, or ecological attributes that were shaped by previous exposure to specific conditions (Schweiger et al. 2019). These retained attributes can greatly affect how communities cope with novel disturbances such that communities that previously experienced disturbances are often more resilient towards novel disturbances (Bengtsson et al. 2003, Johnstone et al. 2016, Hughes et al. 2019). In this study, we focus on an ecological, community-level component of ecological memory, i.e. how species composition, which was shaped by past experiences, influences the response of the community to a novel disturbance.

Coastal areas are among the most productive ecosystems on the planet and have great ecological and economic value (Heip et al. 1995, Harley et al. 2006). Intertidal mudflats harbor a multitude of different species from all domains of life. Microalgae are the main primary producers fueling these diverse benthic food webs (Markert et al. 2013, Rigolet et al. 2014). Benthic microalgae contribute up to 50% of total primary production in some intertidal areas where they can form extensive biofilms on surface sediments (Underwood and Kromkamp 1999, Decho 2000, Stal 2003, Kromkamp et al. 2006). Benthic microalgae biomass and diversity is regulated by many different factors, among them resource availability and grazing (Underwood and Kromkamp 1999, Weerman et al. 2011a, 2011b). The presence of ecosystem engineers such as mussels or oysters also greatly influences benthic microalgae biomass and species composition (Donadi et al. 2013a, Engel et al. 2017). By creating solid structures on intertidal flats, mussel and oyster beds create clear gradients in hydrodynamic conditions and sediment properties (e.g. sediment grain size and organic matter content), which affect species composition and biomass of many organisms including benthic diatoms (Widdows and Brinsley 2002, Donadi et al. 2013b, van der Zee et al. 2012).

In a microcosm experiment with intertidal benthic diatoms, we tested the importance of ecological memory and access to a regional species pool for the community’s resilience to recurring disturbance events. The diatom communities originated from sites with different histories of hydrodynamic stress. We exposed the diatom communities to mechanical disturbance (physical destruction of biofilm), simulated dispersal between the communities with different origin, and measured their chlorophyll *a* concentration (i.e. biomass) over time.

We hypothesized that: (i) The communities originating from locations with different histories of hydrodynamic disturbance had different species compositions; that (ii) the origin of the species communities from the natural gradient of hydrodynamic disturbance determine their resilience to mechanical disturbance (historically higher levels of hydrodynamic stress correlate with high resilience to experimental disturbance); and that (iii) dispersal, through enabling patch-connectivity, mitigates negative impacts of disturbance in the metacommunities.

## Material and methods

### Study organisms and local conditions

We collected benthic diatom communities from three sites on the mudflat off the coast of Schiermonnikoog island, the Wadden Sea, in October of 2015, immediately before the start of the experiment. The three sites were on a transect spanning from the coast seaward and crossing an intertidal mussel bed. Due to the crossing of the mussel bed, the sites differed in exposure to hydrodynamic stress conditions and consequently sediment characteristics (Table 1). Site 1 was unprotected from hydrodynamic stress and in a sandy area coastward of the mussel bed. Site 2 was seaward of the mussel bed with intermediate protection and muddy sediment. Site 3 (hereafter referred to as “low hydrodynamic stress”) was on a mussel bed, where hydrodynamic stress was reduced and the sediment in the bare patches between mussels was muddy and fine grained.

**Table 1.** Site characteristics of the sample origins. (Origin = hydrodynamic stress exposure at origin, Sed. = sediment type). Erosion was measured with dissolution plasters and is used as a proxy for hydrodynamic forcing. Organic matter content in the sediment was calculated for two different depths: 2 cm (Organic Matter) and 0.2 cm (Shallow OM).

| Site | Coordinates | Origin | Sed. | Erosion (%) | Chlorophyll *a* (µg/g) | Organic Matter (%) | Shallow OM (%) |
| --- | --- | --- | --- | --- | --- | --- | --- |
| 1 | N53.47090,<br>E6.22400 | High | Sand | 5.10 | 21.77 | 0.70 | 1.00 |
| 2 | N53.46686,<br>E6.22469 | Medium | Mud | 3.33 | 259.03 | 2.48 | 11.74 |
| 3 | N53.46776,<br>E6.22462 | Low | Mud | 2.77 | 293.40 | 10.04 | 12.76 |

At each site, we collected the top 0.5 cm surface sediment of an area of 0.5 m^2^ to extract the benthic diatom communities to use in the experiment. Additionally, at each site we took sediment cores (diameter: 26 mm) to measure chlorophyll *a* (three cores of 0.2 cm depth pooled onto a piece of aluminum foil and stored in a sealed plastic bag on ice), organic matter content at two different depths: 2 cm and 0.2 cm depth (i.e. shallow OM); placed into sealed plastic bags and stored on ice), and benthic diatom species composition (core with 2 cm depth; placed in sealed plastic bag and stored on ice). We also measured the level of hydrodynamic disturbance at each site by placing dissolution plasters out for two tidal cycles and measuring the dry weight of the plasters before and after exposure to the tides. We transported all samples in cool boxes back to the laboratory (<24h).

In the laboratory, we extracted the motile benthic diatoms from the large area and from the cores separately by spreading out the sediment and placing two layers of lens cleaning tissue onto the sediment. After 5 h of exposure to light, we collected the top tissue and rinsed the diatoms off into culture bottles with sterile filtered North Sea water. We stored the samples from the large area in the dark at 19°C until the start of the experiment (<4h). We fixed the core samples in Lugol’s iodine and determined species composition with the Utermöhl counting technique (Utermöhl 1958) under an inverted microscope. We freeze-dried the sediment chlorophyll *a* samples, and subsequently measured chlorophyll *a* concentrations using a fluorometer (Trilogy) after acetone extraction (90%, dark, -20°C, 48 h) and methods described by Jeffrey and Humphrey (1975). The organic matter content was determined through Loss on Ignition by burning oven dried organic matter samples (48h, 60°C) in a muffle kiln (4h, 550°C).

### Set-up and sampling

We set up the experiment in a climate room with controlled temperature (19°C) and light levels (10.8 µmol m^-2^ s^-1^ and 14:10 light-dark cycle). We used 60-mL-culture flasks (TPP, filter screw cap) as microcosms for this experiment and 40 mL sterile filtered North Sea water (N:Si:P added for final concentrations of 40:40:2.7 µM) as the medium. We carefully exchanged 20 mL of the medium every third day over the course of the experiment to avoid nutrient limitation.

For our fully factorial experiment, we constructed 54 metacommunities out of 162 local communities. We constructed the metacommunities by connecting three local communities with different origin along the hydrodynamic stress gradient so that each metacommunity contained one local community of high, medium, and low hydrodynamic stress. We adjusted the inocula so that all bottles had similar initial diatom abundances. We applied three different mechanical disturbance levels to the communities: no, intermediate, and frequent. We administered disturbance by scraping the bottom of the culture flask with a cell scraper every fourth day for intermediate and every other day for frequent disturbance levels. The no-disturbance treatment was not subject to scraping. Each local community within a metacommunity was subject to the same disturbance treatment. Half of the local communities were assigned to a dispersal treatment, in which we administered dispersal every other day. To administer dispersal, we first carefully turned the bottles three times to suspend the more loosely attached diatoms into the medium. We then pipetted three mL solution (i.e. medium plus suspended diatoms) out of each of the three bottles per metacommunity into a sterile beaker. In the beaker, we mixed the three local community solutions and returned three mL of this mixture into the respective bottles of the same metacommunity. The communities not subject to dispersal, were treated similarly to the dispersal treatment with the exception that no culture was removed from or added to the bottles. This ensured that the dispersal treatment did not affect boundary layer and nutrient uptake dynamics. The experiment ran for 29 days and we sampled destructively three times (i.e. removed the entire flask from the experiment after two, three, and four weeks of growth). Each treatment combination (including the three sampling times) was replicated three times.

On the three sampling days, we scraped the biofilm off the bottom of the culture flasks and homogenized it in the medium by shaking the flask. We filtered 7 mL of the suspended cultures over GF/F filters to determine chlorophyll *a* concentration of the samples. We measured chlorophyll *a* concentration with a fluorimeter (Trilogy) after extraction with 90% acetone. We calculated regional chlorophyll *a* concentration by summing the separate values from each local community within a metacommunity.

### Statistical analysis

Our fully crossed design included the fixed factors sampling day (three levels: 2, 3, 4), origin (three levels: low hydrodynamic stress, medium hydrodynamic stress, and high hydrodynamic stress), disturbance treatment (three levels: no, intermediate, frequent), and dispersal treatment (two levels: no-dispersal and dispersal). We ran a GLM including all factors and combinations to test the effect of sampling day, origin, disturbance, and dispersal on local chlorophyll *a* concentration. Likewise, on the regional scale, we constructed a model testing the effect of sampling day, disturbance, and dispersal on chlorophyll *a* concentration. Subsequently, we compared treatment levels of the significant main effects (origin and disturbance) with Tukey HSD post-hoc tests. All analysis were done in R v.3.4.1 (R Core Team 2017).

## Results

### Local conditions at origin and initial species composition

As expected, the extraction sites of benthic diatoms varied in their characteristics relating to hydrodynamic stress exposure and thus their sediment properties (Table 1). The unprotected Site 1 (hereafter referred to as “high hydrodynamic stress”) had the highest erosion, but lowest organic matter content and chlorophyll *a* concentration (Table 1). Site 2 with intermediate protection (hereafter referred to as “medium hydrodynamic stress) had intermediate erosion and organic matter content, but high chlorophyll *a* concentration (Table 1). The most protected Site 3 (hereafter referred to as “low hydrodynamic stress”) had the lowest erosion, but the highest organic matter content and chlorophyll *a* concentration (Table 1).

The different communities from different origin along the hydrodynamic stress gradient also had varying benthic diatom species composition (Fig. 1). Site “high hydrodynamic stress” was dominated by several larger *Navicula Sp*., whereas Site “low hydrodynamic stress” had a high relative abundance of the smallest *Navicula Sp* (Fig. 1). Site “medium hydrodynamic stress” was dominated by *Pleurosigma aestuarii*, a very large sigmoidal species (Fig. 1).

**Fig. 1.**
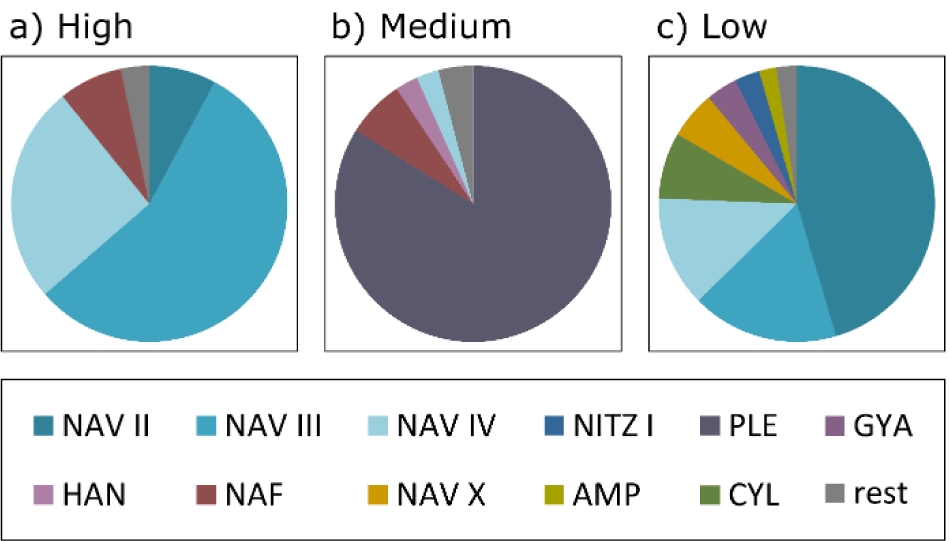
Species composition at different extraction sites (origin). High, medium, and low refer to hydrodynamic stress conditions at the origin (see Table 1). For names and average cell sizes of the different species see Table 2.

### Experimental results

The origin of the diatom communities along the hydrodynamic stress gradient determined the response of the communities to mechanical stress (significant interaction effect between origin and mechanical disturbance: F_4,105_=8.54, p<0.01; Fig. 2). Mechanical disturbance significantly decreased local chlorophyll *a* concentrations in all communities (significant main effect of disturbance: F_2,105_=45.63, p<0.01) with both disturbance treatments (i.e. intermediate (I) and frequent (F) mechanical disturbance) having significantly lower chlorophyll *a* concentrations than the no-disturbance (N) treatment (Tukey HSD: N-I and N-F p<0.01; Fig. 2). However, the higher hydrodynamic stress regime the communities originated from, the more resilient they were to the mechanical disturbance treatment. On average, the mechanical disturbance decreased chlorophyll *a* in the communities with high hydrodynamic stress at origin by 29%, with medium hydrodynamic stress at origin by 44%, and with low hydrodynamic stress at origin by 74% (Tukey HSD origin: high-medium, high-low, and medium-low p<0.01; Fig. 2).

**Table 2.** Abbreviations as used in Fig. 1, names, and average cell size of the most abundant diatom species.

| **ID** | **Species** | **Cell size ($\mu\text{m}^3$)** |
| --- | --- | --- |
| **AMP** | *Amphora Sp. 1* | 787.92 |
| **CYL** | *Cylindrotheca closterium* | 33.16 |
| **GYA** | *Gyrosigma acuminatum* | 1440.38 |
| **HAN** | *Hantzschia Sp. 1* | 612.90 |
| **NAF** | *Navicula forcipata* | 1563.89 |
| **NAV II** | *Navicula Sp. 1* | 49.74 |
| **NAV III** | *Navicula Sp. 2* | 146.92 |
| **NAV IV** | *Navicula Sp. 3* | 730.88 |
| **NAV X** | *Navicula Sp. 4* | 730.88 |
| **NITZ I** | *Nitzschia Sp. 1* | 193.73 |
| **PLE** | *Pleurosigma aestuarii* | 4783.61 |

**Fig. 2.**
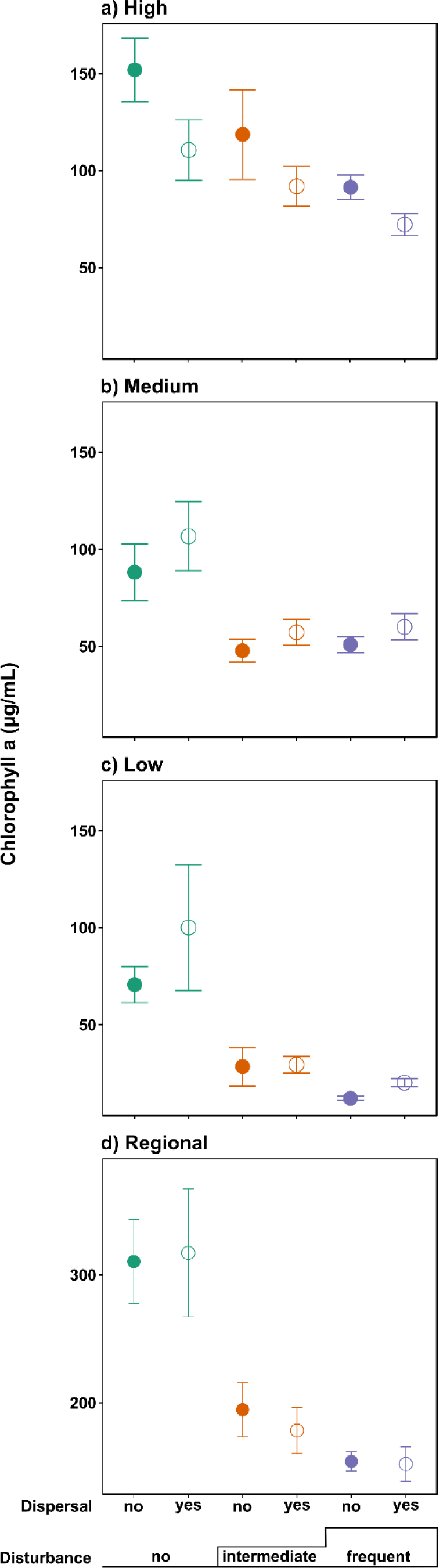
Chlorophyll *a* concentration in the different treatments for each local community with different origin along a hydrodynamic stress gradient and for the regional scale. (a) High hydrodynamic stress at origin, b) medium hydrodynamic stress at origin, c) low hydrodynamic stress at origin, d) regional. Solid circles represent no-dispersal treatments and open circles represent dispersal treatments. No (green), intermediate (orange), and frequent (purple) correspond to the levels of the disturbance treatment in the experiment.

Local chlorophyll a was also significantly affected by interactive effects of origin of the community and dispersal (F_2,105_=5.3, p=0.01). While dispersal decreased chlorophyll *a* concentrations in the local communities originating from highest hydrodynamic stress, it did not affect or slightly increased chlorophyll *a* concentrations in the other communities (origin from medium and low hydrodynamic stress, respectively) (Fig. 2). There was no significant effect of sampling day (Table A1).

Disturbance significantly decreased regional chlorophyll *a* concentration (F_2,35_ = 21.08, p<0.01; N: 313.67±28.29; I: 186.52±13.60; F: 153.20±7.51; Tukey HSD: N-I and N-F p<0.01; Fig. 2), while dispersal and sampling day did not have a significant effect on regional chlorophyll *a* concentration (Fig. 2; Table A2).

## Discussion

Our results demonstrate that the biological properties determined by the origin of the experimental communities along a natural hydrodynamic stress gradient determined their response to new disturbances. The communities with different histories of hydrodynamic stress at origin had different species composition (supporting hypothesis 1). Communities originating from sites that naturally experienced higher levels of hydrodynamic stress had a higher resilience to experimental disturbance than those originating from sites with lower levels of hydrodynamic disturbance (supporting hypothesis 2). However, dispersal did not mitigate negative impacts of disturbance in our experimental metacommunities (rejecting hypothesis 3). Thus, the different communities had different biological properties relating to ecological memory (species composition, diversity, and traits); and the ecological memory of the communities shaped by higher levels of disturbance at origin were more resilient to new disturbances than the communities with lower levels of disturbance at origin.

The variation in species composition at origin (Fig. 1) was likely caused by the differences in local conditions on the intertidal flat including hydrodynamic stress and resulting sediment characteristics (Table 1). Communities previously exposed to higher levels of hydrodynamic stress are probably more resilient to novel disturbances because they are inhabited by species that compensate the disturbance by individual resilience or by rapid growth rates due to previous need for this. Changes in species composition as response to past states is part of the ecological memory of communities. In our study, the ecological memory of the communities originating from the site with high hydrodynamic stress likely contributed to mitigating the negative impact of a novel disturbance on ecosystem functioning. Several other studies also show that ecological memory increases the resilience of communities to novel disturbances in different systems, including terrestrial plants (Johnstone et al. 2016), aquatic plants (Sterk et al. 2016), corals (Hughes et al. 2019), and archaea (Beer et al. 2014). However, it is important to realize that ecological memory is not a universal insurance for resilience, especially considering the projected increase in magnitude and frequency of extreme climate events in the future (Harley et al. 2006, IPCC 2014), which will make disturbance regimes more unpredictable. For example, a recent study by Jacquet and Altermatt (2020) shows that above a certain threshold of past disturbance frequency and intensity, legacy effects can lead to negative effects of past disturbances on present species diversity and ecosystem functioning. The positive effect of ecological memory also depends on species co-tolerance. Only when species’ initial tolerance and tolerance to additional stressors are positively correlated can the impact of an additional stressor be reduced and lead to “stress-induced community tolerance” (Vinebrooke et al. 2004). More research is needed to assess in which cases the positive effect of ecological memory surpasses the negative effect of legacy effects and what other components of ecological memory are important for community resilience.

Previous studies have shown that microalgal species composition and biomass production are dependent on many abiotic and biotic variables and that they are tightly linked to sediment grain size (Cahoon et al. 1999, Thornton et al. 2002, Du et al. 2009). Even though we did not directly measure sediment grain size in this study, visual observations showed that the sandy site with high hydrodynamic stress and low organic matter content had larger grain size. Other studies confirm that locations with high hydrodynamic forcing have larger sediment grain size and lower clay content and therefore are less muddy (de Jong & de Jonge 1995, Thornton et al. 2002, Méléder et al 2007). Smaller species should be able to recover faster after disturbances, because they have higher growth and division rates (Finkel et al. 2010). Therefore, it would be logical to find that communities with smaller species can withstand disturbances better, and thus in our study we would expect to find smaller species in the more highly disturbed sites. We observed the opposite pattern. Species from the high and medium stressed sites were generally larger than those from the low stressed site, independent of sediment grain size at origin. Since in our study the finer grained sediments were in the location of the mussel bed, it is unclear if the sediment determined diatom cell size, or if other factors played into the size selection of species. For example, selective grazing by organisms inhabiting the mussel bed could have influenced benthic diatom species composition and thus possibly led to size discrimination and the presence of predominantly small species in the fine-grained sediment (D’Hondt et al. 2018). In addition, diatoms are generally characterized by high growth rates and maximum nutrient uptake rates, because they are adapted to rapidly responding to nutrient pulses in coastal areas (Litchman et al. 2007). Therefore, the metabolic size scaling might not express in this group of diatoms. Alternatively, this scaling might be masked by local adaptation to hydrodynamic stress in the communities investigated here and thus overridden by ecological memory.

Contrary to expectations, dispersal did not lead to increased chlorophyll *a* concentration on the local nor regional scale, independent of the disturbance level. In the community originating from high stress hydrodynamic conditions, dispersal even decreased chlorophyll *a* compared to the no-dispersal treatment (Fig. 2). Interestingly, the variability of replicates within the dispersal treatment of the no-disturbance communities was high, meaning that the response of these replicates was not uniform. Contrary to the present results, other studies have shown that dispersal indeed “rescues” disturbed communities (Altermatt et al. 2011, Symons & Arnott 2013, Rosset et al. 2017). Through dispersal, communities gain access to a larger species pool on the regional scale so that there should be more species present that have the ideal traits for the novel situation after a disturbance. However, in our experiment, even the intermediate disturbance level seemed too severe, or the dispersal frequency too low, to initiate a rescue effect. Another explanation for our results could be that in our experiment all local communities in a disturbed metacommunity were exposed to the same disturbance level (i.e. the disturbance was a regional event), whereas in nature and in other experiments, the regional species pool oftentimes includes disturbed and undisturbed patches (e.g. Altermatt et al. 2011) so that within the metacommunity dispersal from the undisturbed to disturbed patches can lead to a “rescue effect”. In future experiments, including undisturbed “rescue” patches would be a useful addition to the experimental set-up.

Our experiment shows that initial community composition (specifically relating to ecological memory) largely drives ecosystem functions, despite the presence of other well-known structuring mechanisms, such as dispersal. This exemplifies the important role of species identities for ecosystem functioning in (meta)communities and highlights the crucial need for protecting biodiversity in natural systems.

## References

Altermatt, F., A. Bieger, F. Carrara, A. Rinaldo, and M. Holyoak. 2011. Effects of connectivity and recurrent local disturbances on community structure and population density in experimental metacommunities. PLoS ONE 6:e19525.

Bartholomä, A., A. Kubicki, T. H. Badewien, and B. W. Flemming. 2009. Suspended sediment transport in the German Wadden Sea-seasonal variations and extreme events. Ocean Dynamics 59:213–225.

Beer, K. D., E. J. Wurtmann, N. Pinel, and N. S. Baliga. 2014. Model organisms retain an “ecological memory” of complex ecologically relevant environmental variation. Applied and Environmental Microbiology 80:1821–1831.

Bengtsson, J., P. Angelstam, T. Elmqvist, U. Emanuelsson, C. Folke, M. Ihse, F. Moberg, and M. Nyström. 2003. Reserves, resilience and dynamic landscapes. Ambio 32:389–396.

Beniston, M., D. B. Stephenson, O. B. Christensen, C. A. T. Ferro, C. Frei, S. Goyette, K. Halsnaes, T. Holt, K. Jylhä, B. Koffi, J. Palutikof, R. Schöll, T. Semmler, and K. Woth. 2007. Future extreme events in European climate: An exploration of regional climate model projections. Climatic Change 81:71–95.

Cahoon, L. B., J. E. Nearhoof, and C. L. Tiiton. 1999. Sediment grain size effect on benthic microalgal biomass in shallow aquatic ecosystems. Estuaries 22:735–741.

Cardinale, B. J., J. E. Duffy, A. Gonzalez, D. U. Hooper, C. Perrings, P. Venail, A. Narwani, G.M. Mace, D. Tilman, D. A. Wardle, A. P. Kinzig, G. C. Daily, M. Loreau, J. B. Grace, A. Larigauderie, D. S. Srivastava, and S. Naeem. 2012. Biodiversity loss and its impact on humanity. Nature 486:59–67.

Carrara, F., A. Giometto, M. Seymour, A. Rinaldo, and F. Altermatt. 2015. Experimental evidence for strong stabilizing forces at high functional diversity of aquatic microbial communities. Ecology 96:1340–1350.

Cosentino, B. J., R. L. Schooley, and C. A. Phillips. 2011. Spatial connectivity moderates the effect of predatory fish on salamander metapopulation dynamics. Ecosphere 2(8):1–14.

Decho, A. W. 2000. Microbial biofilms in intertidal systems: An overview. Continental Shelf Research 20:1257–1273.

D’Hondt, A.-S., W. Stock, L. Blommaert, T. Moens, and K. Sabbe. 2018. Nematodes stimulate biomass accumulation in a multispecies diatom biofilm. Marine Environmental Research 140:78–89.

Donadi, S., J. Westra, E. J. Weerman, T. van der Heide, E. M. van der Zee, J. van de Koppel, H. Olff, T. Piersma, H. W. van der Weer, and B. K. Eriksson. 2013a. Non-trophic interactions control benthic producers on intertidal flats. Ecosystems 16:1325–1335.

Donadi, S., T. van der Heide, E. M. van der Zee, J. S. Eklöf, J. van de Koppel, E. J. Weerman, T. Piersma, H. Olff, and B. K. Eriksson. 2013b. Cross-habitat interactions among bivalve species control community structure on intertidal flats. Ecology 94:489–498.

Du, G. Y., M. Son, M. Yun, S. An, and I. K. Chung. 2009. Microphytobenthic biomass and species composition in intertidal flats of the Nakdong River estuary, Korea. Estuarine, Coastal and Shelf Science 82:663–672.

Elmqvist, T., C. Folke, M. Nyström, G. Peterson, J. Bengtsson, B. Walker, and J. Norberg. 2003. Response diversity, ecosystem change, and resilience. Frontiers in Ecology and the Environment 1:488–494.

Engel, F. G., J. Alegria, R. Andriana, S. Donadi, J. B. Gusmao, M. A. van Leeuwe, B. Matthiessen, and B. K. Eriksson. 2017. Mussel beds are power stations on intertidal flats. Estuarine, Coastal and Shelf Science 191:21–27.

Finkel, Z. V., J. Beardall, K. J. Flynn, A. Quigg, T. A. V. Rees, and J. A. Raven. 2010. Phytoplankton in a changing world: Cell size and elemental stoichiometry. Journal of Plankton Research 32:119–137.

Gilpin, M. E., and I. Hanski. 1991. Metapopulation dynamics: Empirical and theoretical investigations. Academic Press, London.

Gonzalez, A., R. M. Germain, D. S. Srivastava, E. Filotas, L. E. Dee, D. Gravel, P. L. Thompson, F. Isbell, S. Wang, S. Kéfi, J. Montoya, Y. R. Zelnik, and M. Loreau, 2020. Scaling‐up biodiversity‐ecosystem functioning research. Ecology Letters 23:757–776.

Harley, C. D. G., A. R. Hughes, K. M. Hultgren, B. G. Miner, C. J. B. Sorte, C. S. Thornber, L. F. Rodriguez, L. Tomanek, and S. L. Williams. 2006. The impacts of climate change in coastal marine systems. Ecology Letters 9:228–241.

Heip, C. H. R., N. K. Goosen, P. M. J. Herman, J. Kromkamp, J. J. Middelburg, and K. Soetaert. 1995. Production and consumption of biological particles in temperate tidal estuaries. Pages 1-149 in Oceanography and Marine Biology: An Annual Review (33).D. Ansell, R. N. Gibson, and M. Barnes, editors. UCL Press.

Holyoak, M., M. A. Leibold, and R. D. Holt. 2005. Metacommunities: Spatial dynamics and ecological communities. University of Chicago Press

Hooper, D. U., F. S. Chapin III, and J. J. Ewel. 2005. Effects of biodiversity on ecosystem functioning: a consensus of current knowledge. Ecological Monographs 75:3–35.

Hughes, T. P., J. T. Kerry, S. R. Connolly, A. H. Baird, C. M. Eakin, S. F. Heron, A. S. Hoey, M. O. Hoogenboom, M. Jacobson, G. Liu, M. S. Pratchett, W. Skirving, and G. Torda. 2019. Ecological memory modifies the cumulative impact of recurrent climate extremes. Nature Climate Change 9:40–43.

IPCC. 2014. Climate Change 2014. Synthesis Report. Contribution of Working Groups I, II and III to the Fifth Assessment Report of the Intergovernmental Panel on Climate Change. R. K. Pachauri and L. A. Meyer, editors. IPCC, Geneva.

Jacquet, C. and F. Altermatt. 2020. The ghost of disturbance past: long-term effects of pulse disturbances on community biomass and composition. Proc. R. Soc. B 287: 20200678.

Jeffrey, S. W., and G. F. Humphrey. 1975. New spectrophotometric equations for determining chlorophylls a, b, c_1_ and c_2_ in higher plants, algae and natural phytoplankton. Biochemie und Physiologie der Pflanzen 167:191–194.

Johnstone, J. F., C. D. Allen, J. F. Franklin, L. E. Frelich, B. J. Harvey, P. E. Higuera, M. C. Mack, R. K. Meentemeyer, M. R. Metz, G. L. W. Perry, T. Schoennagel, and M. G. Turner. 2016. Changing disturbance regimes, ecological memory, and forest resilience. Frontiers in Ecology and the Environment 14:369–378.

de Jong, D. J., and V. N. de Jonge. 1995. Dynamics and distribution of microphytobenthic chlorophyll-a in the Western Scheldt estuary (SW Netherlands). Hydrobiologia 311:21–30.

de Jonge, V. N., and J. E. E. van Beusekom. 1995. Wind- and tide-induced resuspension of sediment and microphytobenthos from tidal flats in the Ems estuary. Limnology and Oceanography 40:776–778.

König, S., M. C. Köhnke, A. Firle, T. Banitz, F. Centler, K. Frank, and M. Thullner. 2019. Disturbance size can be compensated for by spatial fragmentation in soil microbial ecosystems. Frontiers in Ecology and Evolution 7:290.

Kromkamp, J. C., J. F. C. de Brouwer, G. F. Blanchard, R. M. Forster, and V. Créach. 2006. Functioning of microphytobenthos in estuaries. Proceedings of the Colloquium. Royal Netherlands Academy of Arts and Sciences.

Leibold, M. A., M. Holyoak, N. Mouquet, P. Amarasekare, J. M. Chase, M. F. Hoopes, R. D. Holt, J. B. Shurin, R. Law, D. Tilman, M. Loreau, and A. Gonzalez. 2004. The metacommunity concept: A framework for multi-scale community ecology. Ecology Letters 7:601–613.

Leibold, M. A. and J. M. Chase. 2018. Chapter 11: Metacommunity assembly and the functioning of ecosystems. Pages 530–581 in Monographs in Population Biology, Volume 59, Metacommunity Ecology. Princeton University Press, Princeton, USA.

Litchman, E., C. A. Klausmeier, O. M. Schofield, and P. G. Falkowski. 2007. The role of functional traits and trade-offs in structuring phytoplankton communities: Scaling from cellular to ecosystem level. Ecology Letters 10:1170–1181.

Loreau, M., and A. Hector. 2001. Partitioning selection and complementarity in biodiversity experiments. Nature 412:72–76.

Loreau, M., N. Mouquet, and A. Gonzalez. 2003. Biodiversity as spatial insurance in heterogeneous landscapes. Proceedings of the National Academy of Sciences 100:12765–12770.

Markert, A., W. Esser, D. Frank, A. Wehrmann, and K.-M. Exo. 2013. Habitat change by the formation of alien Crassostrea-reefs in the Wadden Sea and its role as feeding sites for waterbirds. Estuarine, Coastal and Shelf Science 131:41–51.

Méléder, V., Y. Rincé, L. Barillé, P. Gaudin, and P. Rosa. 2007. Spatiotemporal changes in microphytobenthos assemblages in a macrotidal flat (Bourgneuf Bay, France). Journal of Phycology 43:1177–1190.

Mori, A. S., T. Furukawa, and T. Sasaki. 2013. Response diversity determines the resilience of ecosystems to environmental change. Biological Reviews 88:349–364.

Mouquet, N and M. Loreau. 2003. Community patterns in source-sink metacommunities. The American Naturalist 162: 544–557.

Ogle, K., J. J. Barber, G. A. Barron-Gafford, L. P. Bentley, J. M. Young, T. E. Huxman, M. E. Loik, and D. T. Tissue. 2015. Quantifying ecological memory in plant and ecosystem processes. Ecology Letters 18:221–235.

Oliver, T. H., M. S. Heard, N. J. B. Isaac, D. B. Roy, D. Procter, F. Eigenbrod, R. Freckleton, A. Hector, C. D. L. Orme, O. L. Petchey, V. Proença, D. Raffaelli, K. B. Suttle, G. M. Mace, B. Martín-López, B. A. Woodcock, and J. M. Bullock. 2015. Biodiversity and Resilience of Ecosystem Functions. Trends in Ecology & Evolution 30:673–684.

Padisak, Judit. 1992. Seasonal succession of phytoplankton in a large shallow lake (Balaton, Hungary) - a dynamic approach to ecological memory, its possible role and mechanisms. Journal of Ecology 80:217–230.

R Core Team. 2017. R: A language and environment for statistical computing.

Rigolet, C., E. Thiébaut, and S. F. Dubois. 2014. Food web structures of subtidal benthic muddy habitats: Evidence of microphytobenthos contribution supported by an engineer species. Marine Ecology Progress Series 500:25–41.

Rosset, V., A. Ruhi, M. T. Bogan, and T. Datry. 2017. Do lentic and lotic communities respond similarly to drying?? Ecosphere 8:e01809.

Schweiger, A. H., I. Boulangeat, T. Conradi, M. Davis, and J.-C. Svenning. 2019. The importance of ecological memory for trophic rewilding as an ecosystem restoration approach. Biological Reviews 94:1–15.

Stal, L. J. 2003. Microphytobenthos, their extracellular polymerics, and the morphogenesis of intertidal sediments. Geomicrobiology Journal 20:463–478.

Sterk, M., G. Gort, H. De Lange, W. Ozinga, M. Sanders, K. Van Looy, and A. Van Teeffelen. 2016. Plant trait composition as an indicator for the ecological memory of rehabilitated floodplains. Basic and Applied Ecology 17:479–488.

Symons, C. C., and S. E. Arnott. 2013. Regional zooplankton dispersal provides spatial insurance for ecosystem function. Global Change Biology 19:1610–1619.

Thornton, D. C. O., L. F. Dong, G. J. C. Underwood, and D. B. Nedwell. 2002. Factors affecting microphytobenthic biomass, species composition and production in the Colne Estuary (UK). Aquatic Microbial Ecology 27:285–300.

Underwood, G. J. C., and J. Kromkamp. 1999. Primary production by phytoplankton and microphytobenthos in estuaries. Advances in Ecological Research 29:93–153.

Utermöhl, H. 1958. Zur Vervollkommnung der quantitativen Phytoplankton Methodik. Mitteilungen der Internationalen Vereinigung der Theoretischen und Angewandten Limnologie 9:263–272.

Vinebrooke, R. D., K. L. Cottingham, J. Norberg, M. Scheffer, S. I. Dodson, S. C. Maberly, and U. Sommer. 2004. Impacts of multiple stressors on biodiversity and ecosystem functioning: the role of species co‐tolerance. Oikos 104: 451–457.

Weerman, E. J., P. M. J. Herman, and J. van de Koppel. 2011a. Top-down control inhibits spatial self-organization of a patterned landscape. Ecology 92:487–495.

Weerman, E. J., P. M. J Herman, and J. van de Koppel. 2011b. Macrobenthos abundance and distribution on a spatially patterned intertidal flat. Marine Ecology Progress Series 440:95–103.

Widdows, J., and M. Brinsley. 2002. Impact of biotic and abiotic processes on sediment dynamics and the consequences to the structure and functioning of the intertidal zone. Journal of Sea Research 48:143–156.

Wilson, D. S. 1992. Complex Interactions in Metacommunities, with Implications for Biodiversity and Higher Levels of Selection. Ecology 73:1984–2000.

van der Zee, E. M., T. van der Heide, S. Donadi, J. S. Eklöf, B. K. Eriksson, H. Olff, H. W. van der Veer, and T. Piersma. 2012. Spatially extended habitat modification by intertidal reef-building bivalves has implications for consumer-resource interactions. Ecosystems 15:664–673.

