## Supplementary material for "Ecological memory mitigates negative impacts of disturbance on biomass production in benthic diatom metacommunities": Table A1.

Table A1. ANOVA results for local chlorophyll *a* (log10(chla) ~ day*origin*disturbance*dispersal). Significant results ($p < 0.05$) are displayed in bold.

| | Df | Sum Sq | Mean Sq | F value | Pr(>F) |
|---|---|---|---|---|---|
| day | 2 | 0.24 | 0.12 | 3.05 | 0.05 |
| **disturbance** | **2** | **3.50** | **1.75** | **45.63** | **0.00** |
| dispersal | 1 | 0.04 | 0.04 | 1.01 | 0.32 |
| **origin** | **2** | **7.91** | **3.96** | **103.03** | **0.00** |
| day:disturbance | 4 | 0.23 | 0.06 | 1.50 | 0.21 |
| day:dispersal | 2 | 0.16 | 0.08 | 2.03 | 0.14 |
| disturbance:dispersal | 2 | 0.05 | 0.03 | 0.64 | 0.53 |
| day:local | 4 | 0.29 | 0.07 | 1.90 | 0.12 |
| **disturbance:origin** | **4** | **1.31** | **0.33** | **8.54** | **0.00** |
| **dispersal:origin** | **2** | **0.41** | **0.20** | **5.31** | **0.01** |
| day:disturbance:dispersal | 4 | 0.10 | 0.03 | 0.66 | 0.62 |
| day:disturbance:origin | 8 | 0.37 | 0.05 | 1.21 | 0.30 |
| day:dispersal:origin | 4 | 0.12 | 0.03 | 0.78 | 0.54 |
| disturbance:dispersal:origin | 4 | 0.06 | 0.02 | 0.40 | 0.81 |
| day:disturbance:dispersal:origin | 8 | 0.17 | 0.02 | 0.55 | 0.82 |
| residuals | 105 | 4.03 | 0.04 | | |
