## Supplementary material for "Ecological memory mitigates negative impacts of disturbance on biomass production in benthic diatom metacommunities": Table A2.

Table A2. ANOVA results for regional chlorophyll *a* (log10(chla) ~ day*disturbance*dispersal). Significant results ($p < 0.05$) are displayed in bold.

| | Df | Sum Sq | Mean Sq | F value | Pr(>F) |
|---|---|---|---|---|---|
| day | 2 | 0.06 | 0.03 | 1.72 | 0.19 |
| **disturbance** | **2** | **0.79** | **0.39** | **21.08** | **0.00** |
| dispersal | 1 | 0.01 | 0.01 | 0.35 | 0.56 |
| day:disturbance | 4 | 0.09 | 0.02 | 1.20 | 0.33 |
| day:dispersal | 2 | 0.08 | 0.04 | 2.24 | 0.12 |
| disturbance:dispersal | 2 | 0.00 | 0.00 | 0.03 | 0.97 |
| day:disturbance:dispersal | 4 | 0.03 | 0.01 | 0.45 | 0.77 |
| residuals | 35 | 0.65 | 0.02 | | |
